# A modular flow-through system for continuous physiological measurements of aquatic holobionts

**DOI:** 10.64898/2026.11.22.768396

**Authors:** Hunt Jones, Claudia Pogoreutz, Christine Ferrier-Pagès, Renaud Grover, Anders Meibom, Nils Rädecker

## Abstract

1. Physiological processes, such as primary productivity, respiration, and nutrient fluxes, are pivotal to marine biogeochemical cycles and the functioning of coastal and estuarine ecosystems. However, conventional laboratory methods to quantify these processes cannot mimic the rapidly varying conditions that characterize coastal environments.
2. We report the development of a modular flow-through incubation system that allows for continuous monitoring of physiological processes in benthic holobionts with a temporal resolution of minutes, over the course of days to weeks. The system consists of flow-through chambers with sensors placed at the inflow, outflow, and onto the chambers themselves, permitting parallel monitoring of physiological activity using, e.g., O_2_ sensing and PAM fluorometry, as well as isotopic labeling and time-point sampling for tissue and omics analyses. The entire system fits into a programmable incubator that establishes highly controlled, yet dynamic environmental conditions.
3. Using the photosymbiotic sea anemone *Exaiptasia diaphana*, we show that the system can accurately quantify metrics of holobiont physiology under variable environmental conditions with high resolution. The system can monitor temporal dynamics of net productivity and respiration rates, and detect differences between light and dark respiration, allowing robust estimates of productivity. For example, daytime net O_2_ production is five times higher with our system than in closed chambers. Importantly, the system may also minimize experimental bias by allowing for incubations with environmentally relevant nutrient concentrations in isotopic labelling experiments, which has not been done with conventional closed-system incubations.
4. The flow-through system thus constitutes a novel experimental platform adaptable to a broad range of biological questions. By allowing continuous monitoring over extended periods of time, it will improve our capacity to investigate the effects of future global change scenarios on benthic organisms.

## Introduction

Benthic invertebrates are important drivers of marine biogeochemical cycles and are foundational to coastal and estuarine ecosystem functioning (Sivadas et al., 2020). However, anthropogenic change, including eutrophication, sedimentation, and ocean warming (Hughes et al., 2017; Kennish, 2022; Straub et al., 2022) have caused widespread ecosystem destruction and restructuring over the past decades (Hoegh-Guldberg, 1999; Kelly et al., 2012; Lesser et al., 2016). Understanding effects of environmental change on biogeochemical processes and physiological performance of marine invertebrates is therefore critical to assess current and future ecosystem functioning. However, temporal variability inherent to coastal systems remains experimentally challenging. Pronounced fluctuations in environmental conditions, ranging from temperature to nutrient availability, have direct effects on the physiology, performance, and role of benthos in coastal ecosystems (Safaie et al., 2018; Saurel et al., 2007; Sawall et al., 2022). Such environmental fluctuations may exert positive or negative effects from individual to community levels. For example, prolonged exposure to high diurnal temperature variability may improve thermotolerance in tropical corals (Safaie et al., 2018; Thomas et al., 2018). In contrast, less predictable perturbations like intense rainfall, run-off, or internal waves cause rapid changes to key environmental factors, such as turbidity, temperature, and water chemistry, often exerting adverse effects on native communities (Lan et al., 2024). Thus, environmental variability on different spatiotemporal scales has diverse and consequential effects. However, most existing methods to evaluate physiological responses of benthic organisms cannot adequately capture this variability (Hardison et al., 2026).

Conventional methods used to study benthic physiological processes, such as calcification, metabolism, and nutrient assimilation, are often based on closed system incubations with controlled conditions (Camp et al., 2015; Liu et al., 2020; Lyndby et al., 2020; Rädecker et al., 2017; Ross et al., 2022; Sawall et al., 2022). To avoid oversaturation or depletion of nutrients or O_2_ during incubations, this approach is limited to short periods with static conditions. While such incubations efficiently quantify physiological performance at holobiont (a host organism and its associated microbes) scale, they generally cannot address complex environmental variability typical of shallow marine habitats. For example, conventional respirometry commonly assumes constant metabolic rates over diurnal cycles to estimate energy budgets (Camp et al., 2015; Hudspith et al., 2022; Sawall et al., 2022). Depending on organismal metabolism and chamber volume, static incubations last from minutes to a few hours, during which metabolism is estimated according to steady rates of change in [O_2_] within incubation chambers (Hudspith et al., 2022; Kühl et al., 1996; Owen et al., 2021; Sawall et al., 2025), making dynamic organismal responses difficult to account for. In contrast, metrics of O_2_ flux with high temporal and spatial resolution have been achieved using microsensing (Kühl et al., 1995; Zhang et al., 2025), however, tissue heterogeneity at small spatial scales renders scaling results to the holobiont scale difficult.

Finally, isotopic labelling experiments using closed-system incubations, coupled with mass spectrometry, have proven a powerful approach to quantify uptake, incorporation, and release of organic and inorganic compounds in marine organisms. When applied to complex holobionts, these approaches permit resolving host/symbiont metabolic interactions (Ferrier-Pagès & Leal, 2019). However, isotopic labelling studies commonly utilize high (often environmentally unrealistic) concentrations of labeled substrates to avoid depletion during incubations and only provide snapshots from which uptake rates are extrapolated (Achlatis et al., 2018; Freeman et al., 2013; Hudspith et al., 2021; Kopp et al., 2015; Pernice et al., 2012; Rädecker et al., 2023).

Thus, existing methodologies, while highly complementary, have shortcomings that limit our ability to assess effects of inherently dynamic conditions on benthos of shallow-water ecosystems. Here, using the photosymbiotic sea anemone *Exaiptasia diaphana* as a model system, we present a flow-through experimental system capable of overcoming these methodological challenges and provide novel insights into the physiology of benthic organisms in variable environments. Our flow-through system builds upon past methods used to estimate metabolism (Liu et al., 2020; Rädecker et al., 2017; Ross et al., 2022; Sawall et al., 2025) by providing high temporal resolution monitoring of biological activities under controlled, dynamic experimental conditions for timescales extending up to two weeks. The modular design allows incorporation of multiple sensors, rendering the system adaptable to diverse research interests.

## Methods

### Flow-through system

The experimental system (Figure 1) is a modular flow-through design, permitting simultaneous monitoring of multiple incubation chambers. A LabMate4 peristaltic pump with four channels (Acmer; Hebei, China) maintains constant flow of seawater from an aerated reservoir through transparent incubation chambers. Luer-locks connect system components, facilitating easy assembly and modularity, allowing exchange of sensors or incorporation of multiple sensors in parallel.

**Figure 1:**
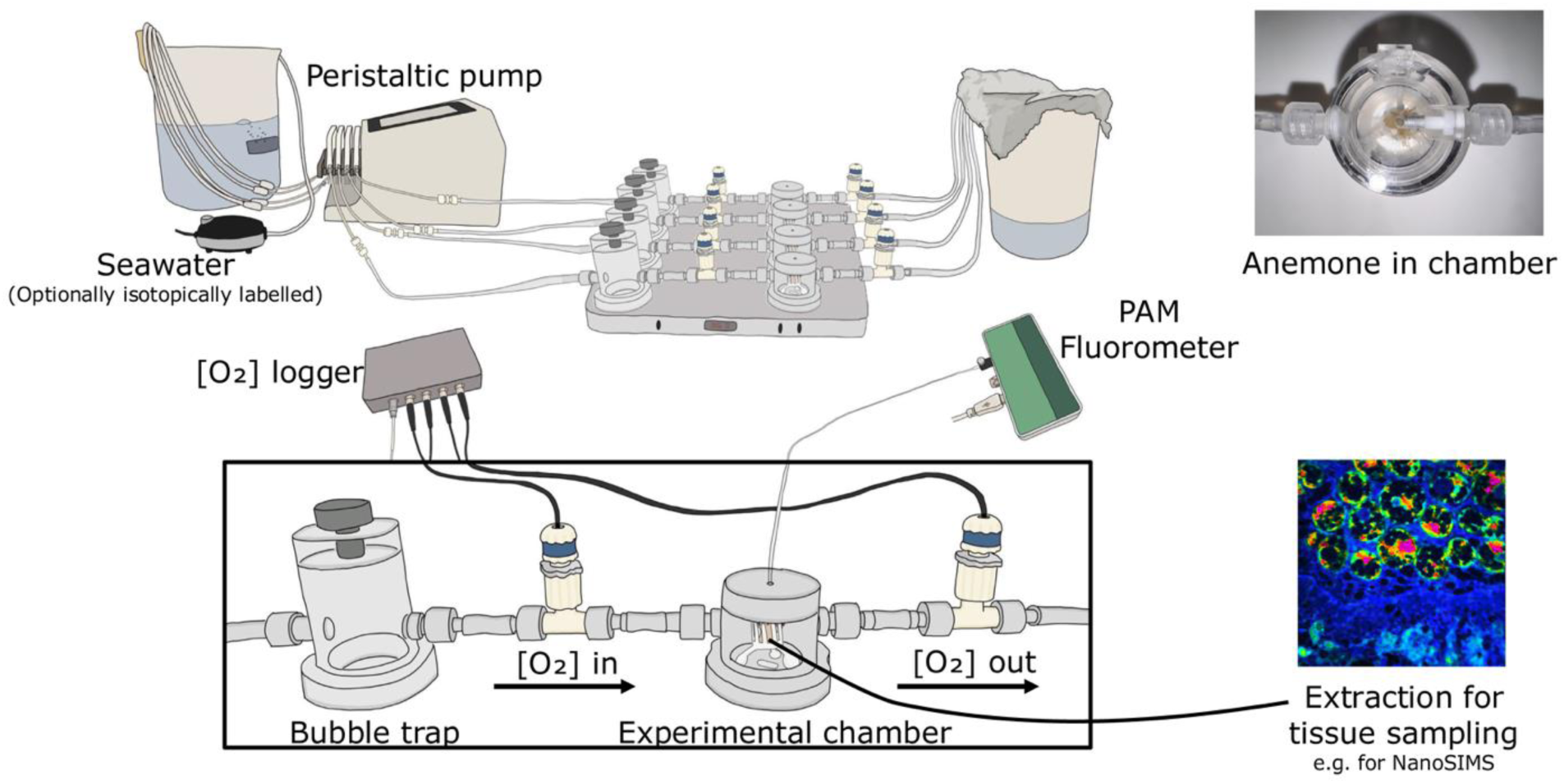
Schematic of flow-through system for simultaneous real-time biological rate measurements using four flow-through chambers in parallel. Monitoring of biological activity is performed with sensors placed at inflow and outflow of each chamber. In the current study, one chamber contained only seawater to account for pelagic background metabolism, while others each contained one animal. Design files for recreation of chambers and 3D printed stages are available in supplemental material (**Files S1, S2, S3**).

Prior to incubations, study organisms are settled onto small stages (File S1) that are then fixed inside incubation chambers (File S2) on mounting points at the chamber’s base. The chamber design allows for a magnetic stir bar below the stage, ensuring constant mixing within the chamber without disturbing study organisms.

Here, Firesting OXFLOW O_2_ sensors (Pyroscience; Aachen Germany) are placed at the inflow and outflow of each chamber (Figure 1), providing continuous monitoring of [O_2_]. To prevent potential air bubbles from interfering with real-time measurements, bubble traps (File S3) are placed before the inflow O_2_ sensors (Figure 1). In addition to O_2_ sensors, a pulse-amplitude-modulation (PAM) fluorometer can be installed atop each chamber, providing photophysiological metrics. Other sensors (e.g., pH and temperature sensor spots) can be connected via optical ports on the side of the chamber or connected upstream/downstream of the chamber to monitor physicochemical parameters. At the end of experimentation, samples are easily extracted for further manipulation, such as processing for omics, isotope measurements, or advanced imaging.

The flow-through system provides a constant supply of seawater (here 1 mL min^-1^) with highly controlled properties, such as specific nutrient concentrations, isotope tracers, or ecotoxin dosing. Here, the entire system is housed within a programmable incubator (CU-22L, Percival, USA), allowing for dynamic light and temperature regulation.

To illustrate the capabilities of our system, we performed a series of experiments to: (1) compare productivity and photochemistry under contrasting light regimes; (2) estimate daytime respiration; (3) compare net productivity and respiration in flow-through and conventional closed chamber incubations; (4) monitor long-term (2 weeks) trends of organismal physiological performance within the system; and (5) evaluate patterns of nutrient assimilation following labeling experiments with low and elevated nutrient concentrations, respectively.

For these experiments, we used the photosymbiotic sea anemone model *Exaiptasia diaphana* strain CC7, hosting its native algal symbiont community dominated by *Symbiodinium linucheae* strain SSA01. A permanent laboratory culture of *E. diaphana* was maintained in light and temperature-controlled incubators (CU-22L, Percival LifeScience) at constant 25°C on a 12h:12h light:dark schedule with constant irradiance of 80 μmol photoquanta m^-2^ sec^-1^ during the photoperiod, and fed ad libitum weekly with freshly hatched *Artemia salina* nauplii.

Preceding each experiment, *E. diaphana* were starved for one week. One day prior to experimentation, individuals were settled onto experimental stages, affixed into chambers and placed within the flow-through system. Flow was maintained at 1 mL min^-1^ (confirmed by pump calibration preceding each experiment) throughout all experiments, providing a chamber residence time of 11.3 minutes for the chamber volume (11.3 mL). *E. diaphana* were given a minimum of 6 hours to acclimate within the system before commencing data collection at midnight.

### Experiment 1 - Effects of irradiance regimes on net productivity and photochemical quantum yield

We first demonstrated capability of continuous net productivity (NP) monitoring over multiple days under static irradiance (SI) and dynamic irradiance (DI) regimes, where NP is the net O_2_ flux resulting from O_2_ production via photosynthesis and O_2_ consumption via respiration (R) in the holobiont.

To investigate effects of variable irradiance on NP throughout the day, *E. diaphana* (n = 4) were incubated for 24 hours under SI followed by 24 hours under DI, with the SI regime identical to general animal husbandry conditions. In contrast, the DI regime consisted of a 12h:12h light:dark cycle with daytime irradiance mimicking a natural irradiance curve, ranging from 32 to 109 μmol photoquanta m^-2^ sec^-1^ with a Gaussian shape during daytime (see Supplemental methods). Importantly, SI and DI treatments delivered equal integrated net photon flux (approximately 3.5 mol photoquanta m^-2^) over the 12-hour light period.

After 48 hours of continuous NP monitoring, total O_2_ production under each treatment was calculated by integrating NP throughout each 12-hour photoperiod for each individual, and mean total daily O_2_ production was compared between irradiance regimes.

Here, NP was estimated using measured net O_2_ flux of each chamber, which was determined according to Equation 1:

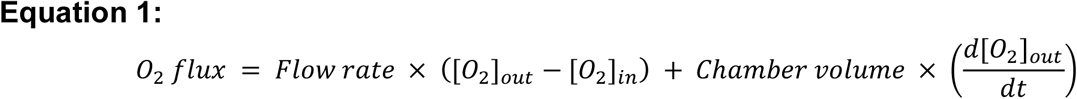

where [O_2_] (in μmol O_2_ liter^-1^) is measured every second by O_2_ sensors (calibrated using aerated seawater preceding each experiment) at inflow and outflow of each chamber. d[O_2_]/dt is the rate of change in [O_2_] calculated at each point in time (t) using a moving window of 20 seconds (d[O_2_]_in_/dt is excluded from Equation 1 because it is in practice negligible). NP was then estimated according to the O_2_ flux of chambers containing holobionts, corrected against O_2_ flux of the negative control. The first term in Equation 1 thus represents NP from steady-state metabolism, while the second represents dynamic changes in NP induced by changing environmental conditions that affect GP and/or R. NP was then normalized to host protein content (see Supplemental methods for details) for all individuals and is reported in standardized units of μmol O_2_ s^-1^ g host protein^-1^.

A complementary experiment was conducted to compare effective (Y(II)) and maximum (dark-adapted; Fv/Fm) quantum yield for *E. diaphana* holobionts (n = 4 per treatment), obtained with PAM fluorometry throughout 24 hours (Genty et al., 1989; Schreiber et al., 1995). PAM measurements were made once per hour using a MINI-PAM-II (blue version) connected to a computer with WinControl-3 software (Walz Photosynthesis Instruments), administering saturating pulses of light at the start of each hour. Measurements in daytime provided metrics of Y(II), while those at night provided Fv/Fm, as Y(II) = Fv/Fm during darkness.

### Experiment 2 - Estimating daytime R

We observed transitions from productivity to respiration within 6 minutes of light/dark transitions at the end of each day in Experiment 1 (Figure 2). Thus, we designed an experiment to exploit the responsiveness of our system to changes in organismal metabolism, and estimate daytime R by interrupting the photoperiod during midday. Here, *E. diaphana* (n = 5) were placed in the flow-through system for 48 hours of continuous NP monitoring. Experimental conditions were identical to those of the SI regime described above, except that we introduced short 6-minute periods of darkness at 11:00 AM, noon, and 1:00 PM (when NP was most stable during prior experiments). During these periods, photosynthesis stopped, resulting in a rapid decrease in outflow [O_2_] that allows estimation of R using Equation 1. Daytime R was then estimated according to the maximal R measured during these periods, and normalized to host protein content.

**Figure 2:**
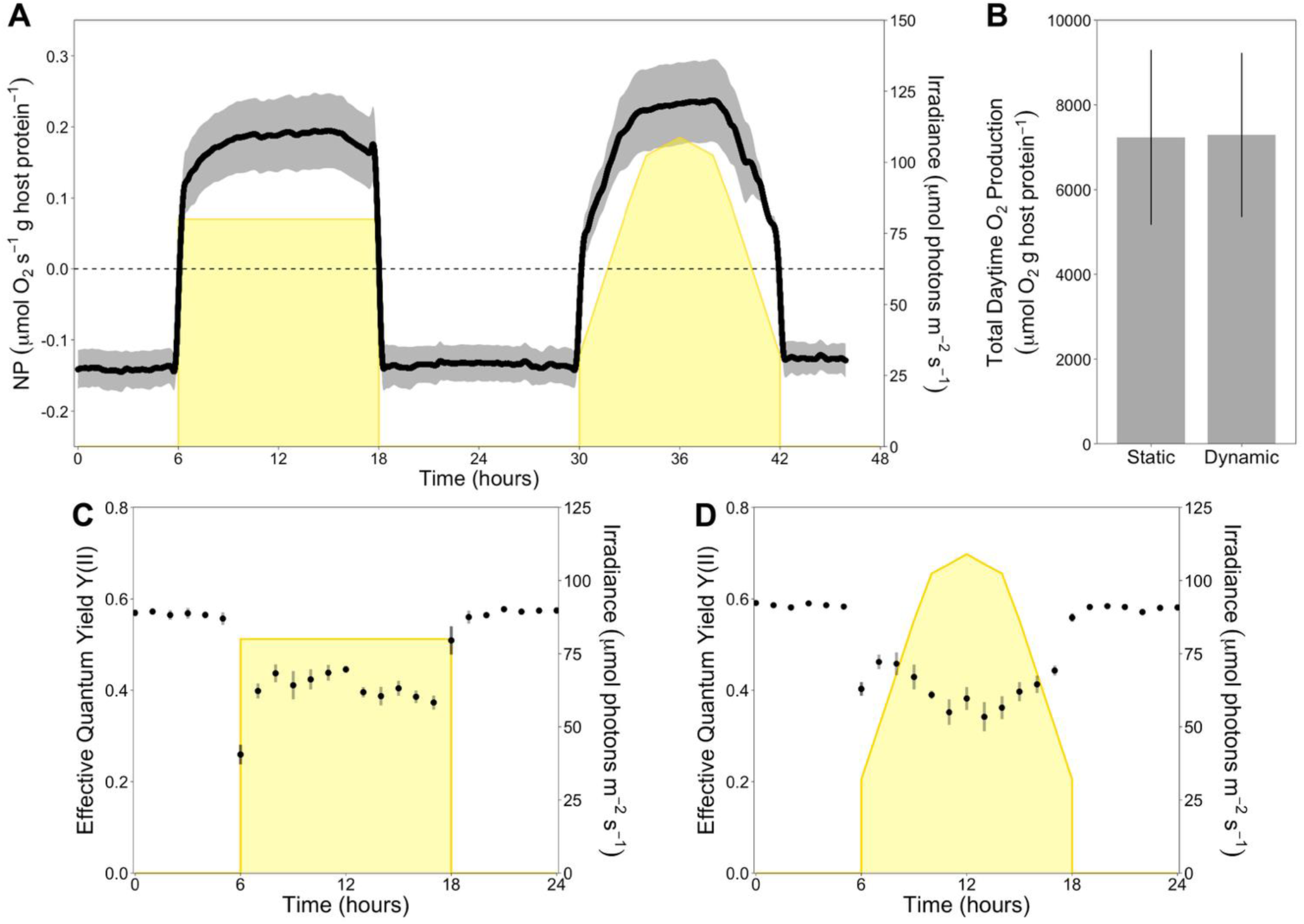
(A) NP of photosymbiotic *Exaiptasia diaphana* CC7 holobionts (n = 4) monitored continuously in the flow-through system over 48h, with first a SI- and then DI photoperiod. (B) Comparison of mean (± SE) daytime O2 production between irradiance regimes. (C and D) Y(II) (effective quantum yield) monitored for 24h under both static and dynamic irradiance (n = 4 for each treatment). In darkness, Y(II) is equal to Fv/Fm (maximum quantum yield). In panels A, C, and D, black lines and points surrounded by grey shading represent the mean ± SE for each measurement over time. Yellow areas represent irradiance (SI vs DI) during each photoperiod.

### Experiment 3 - Comparisons between the flow-through system and conventional methodology

To provide a baseline to which the flow-through system could be compared, experiments were also conducted using conventional closed-system incubation chambers to estimate NP and R. The airtight chambers were nearly identical to those used in flow-through experiments, but without in-and outflow connections, and a PyroScience FireSting O_2_ sensor spot placed within each chamber for continuous monitoring of [O_2_]. While constantly submersed, *E. diaphana* (n = 5) were settled onto stages for several hours, and the stages were then placed in the airtight chambers with magnetic stirring at 160 rpm, ensuring constant mixing. The animals were acclimated for 30 min in darkness before [O_2_] was measured every two seconds in light (80 μmol photoquanta m^-2^ sec^-1^) for one hour. Then, [O_2_] was measured every two seconds in darkness for one hour to estimate R. Estimates of NP and R were calculated from linear fits of [O_2_]/t within each chamber (R statistical software v4.5.0; R Core Team 2025). We identified periods of fifteen minutes in both light and darkness that exhibited the highest linearity based on R^2^ values (> 0.98). During these 15 minute periods, the slope from linear fit models (d[O_2_]/dt) was used to estimate NP and R, respectively, according to:

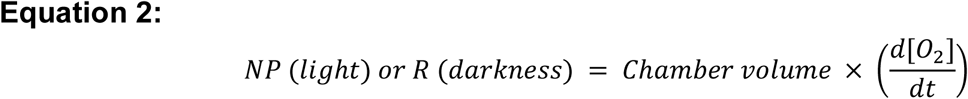

NP and R were corrected for pelagic background metabolism measured simultaneously in a control chamber containing only seawater, then normalized to host protein content. Normalized NP and R were then extrapolated as constant rates over 12h in light (NP) and in darkness (R), respectively, to obtain an estimate of total daily O_2_ production, which was compared to results from the flow-through system.

Estimates of total O_2_ production during the photoperiod, total O_2_ consumption during the night, total O_2_ production over 24 hours, and the ratio of O_2_ production during the photoperiod to O_2_ consumption at night (a proxy of P/R) were made for both closed and flow-through systems.

### Experiment 4 - Long-term NP monitoring in flow-through

As a final test of the flow-through system, we held an individual *E. diaphana* within the system for two weeks to evaluate whether prolonged incubations induce mortality or other significant impacts on physiological performance. The anemone was transferred into the flow-through system and held for two weeks with DI photoperiods and continuous NP monitoring. An additional chamber was connected to provide a negative control. During two weeks, flow was maintained at constant one mL min^-1^ and [O_2_]_out_ and [O_2_]_in_ were measured every one second.

### Experiment 5 - Isotopic labeling experiments

The constant supply of new seawater in the flow-through system effectively removes the risk of nutrient depletion during incubations, permitting investigation of how nutrient concentrations affect assimilation patterns in stable isotope labeling experiments. We compared ^15^N stable isotope assimilation in *E. diaphana* under high (10 μM) and low (1 μM) concentration of ^15^NH_4_^+^ using isotope ratio mass spectrometry (IRMS) and Nanoscale Secondary Ion Mass Spectrometry (NanoSIMS). Incubations lasted 24 hours (one 12-hour SI photoperiod and 12 hours darkness) with a flow rate of 1 mL min^-1^. *E. diaphana* (n = 4) were placed into the system and acclimated in unlabelled artificial seawater overnight before the flow was switched to labelled seawater the next morning, just before the photoperiod began.

For bulk IRMS analysis, artificial seawater was prepared according to (Harrison et al., 1980) containing either 10 μM or 1 μM of ^15^NH ^+^. At the end of the experiment, isotopically labeled holobionts (n = 4 per treatment plus 2 unlabeled controls) were collected and immediately stored at-20°C to await further processing.

For NanoSIMS analysis, we prepared artificial seawater as described above, with treatments of either 10 μM or 1 μM concentration of ^15^NH ^+^. After 24 hours of labelling, holobionts (n = 3 in the 10 μM treatment and 4 in the 1 μM treatment; plus 2 unlabeled controls) were collected and immediately fixed in a solution of 2.5% glutaraldehyde and 1% paraformaldehyde in PBS with 9% weight/volume sucrose for 24 hours before being stored in in PBS with 9% weight/volume sucrose at 4°C until further processing.

### Stable isotope analysis of bulk tissue

Frozen samples were thawed at room temperature, homogenized in 500 μL milliQ H_2_O and centrifuged at 3,000 × g for 3 minutes to separate host and symbiont cell fractions. The supernatant containing host material was removed and pipetted into a glass vial before being refrozen. The algal pellet was then resuspended in 500 μL milliQ H_2_O and pipetted into a glass vial before being refrozen.

Frozen samples were lyophilized for approximately 32 hours using a Christ Beta 2-8 LD-plus lyophilizer (Martin Christ; Osterode am Harz, Germany). Dried material was then rehomogenized within each vial using a clean micropestle and weighed into small tin capsules using a microbalance to obtain two technical replicates of approximately 350 μg per sample for analysis of ^15^N enrichment. The ^15^N/^14^N ratios for each sample were obtained using an isotope ratio mass spectrometer coupled to an elemental analyzer and converted to atom percent excess (APE) according to:

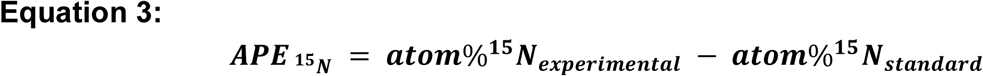

where ***atom***%**^15^*N*** was calculated by:

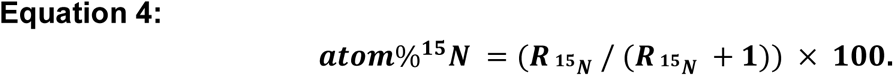

### Correlative scanning electron microscopy and NanoSIMS imaging

Samples fixed for NanoSIMS analysis were processed following the methods of (Rädecker et al., 2023). In brief, dissected tentacles were washed, stained in 1% OsO_4_, dehydrated in an ethanol series, then infiltrated in a series of increasing concentrations of SPURR resin in acetone, before being left to infiltrate overnight in 100% SPURR, then transferred into fresh resin for polymerization at 65°C. Semi-thin (200 nm) sections were then taken using an ultramicrotome (Leica Microsystems; Saterland, Germany), and placed onto silicon wafers. SEM imaging of these sections was done with a Gemini 500 SEM instrument (Carl Zeiss; Oberkochen, Germany) at the Center for Advanced Surface Analysis at the University of Lausanne using backscatter detection and an accelerating voltage of 3 kV. SEM imaging served to identify areas suitable for subsequent NanoSIMS imaging and identify cell types and tissue layers within those areas.

The isotopic composition of sample sections was analyzed using a NanoSIMS 50L instrument (Hoppe et al., 2013). Samples were sputter-coated with gold (∼15 nm) prior to being placed in the NanoSIMS. Mass images of 45×45 µm^2^ areas of the sample surface were generated, and then analyzed using the OpenMIMS plugin for ImageJ (Poczatek et al., 2012), following the settings detailed by (Rädecker et al., 2023). Supported by SEM images, 8 sample areas were analyzed for each biological replicate. The final ^12^C^14^N^-^ images were used to distinguish symbiotic algal cells from adjacent host tissue and manually define individual regions of interest (ROI). For each ROI, the ^12^C^15^N^-^/^12^C^14^N^-^ ratio was obtained and isotopic enrichment values were expressed in APE. Enrichment in ^15^N was calculated according to Equation 3 and Equation 4. Unlabelled samples were analyzed to obtain standard values and ***atom***% **^15^*N_standard_*** represents the mean value obtained from comparable ROIs of these unlabelled control samples.

### Statistical analysis

#### Effects of irradiance regime on NP and photophysiolog

The average of the total O_2_ production over a 12-hour photoperiod under SI and DI was compared using Student’s paired t-test.

Linear and polynomial regression analyses were used to investigate trends in Y(II) throughout the photoperiod and Fv/Fm during the night, separately. The first morning measurement was excluded from these analyses, as illumination of incubator lights just before PAM measurements resulted in high fluorescence preceding the saturating pulse of light, driving Y(II) down without causing photoinhibition (Figure 2C, D) (Schreiber et al., 1995). We next tested for differences in Fv/Fm at night among treatments using a linear mixed model with time and treatment as fixed effects, and sample as a random effect.

#### Differences in day-and nighttime respiration

Day-and nighttime respiration were compared by aggregating measurements for each individual taken during the light phase and dark phase, respectively, and then conducting a paired Student’s t-test.

#### Comparison of NP in closed and flow-through systems

Estimates of total O_2_ production during a 12-hour photoperiod, total O_2_ consumption during a 12-hour night, and total O_2_ production over a 24-hour day/night cycle were compared between flow-through and closed incubation systems using Student’s t-tests. Estimates of total O_2_ production during a 12-hour photoperiod were reciprocal transformed to satisfy the assumption of normality; otherwise no test assumptions were violated in any comparison. Equal variance among groups was confirmed for each comparison via Levene’s test.

#### Long-term monitoring of NP in flow-through

With only one individual, no statistical tests are presented here, but summary statistics are included, describing physiological performance of *E. diaphana* over two weeks with continuous O_2_ monitoring.

#### Host and symbiont nutrient uptake affected by NH ^+^ concentration

Enrichment in ^15^N (in APE) of bulk tissue measured via IRMS was compared among treatments and symbiotic partner identity using a two-way ANOVA. Additionally, we compared ^15^N enrichment (in APE) measured using NanoSIMS among treatments and symbiont partner identity using a linear mixed model LMM, generated with the R package lmer4 (Bates et al., 2015), with ^15^N enrichment as the response variable, treatment and host/symbiont identity as fixed effects, and sample as a random effect.

All statistical analyses were conducted using R statistical software (v4.5.0; R Core Team 2025). All data are presented as means ± standard error (SE).

## Results

The flow-through system provided reliable, continuous monitoring of physiological performance of *E. diaphana* holobionts across five complementary experiments, demonstrating its efficacy in resolving temporal dynamics of holobiont physiology in dynamic environments.

### Experiment 1 - Effects of light regime on NP and photosynthetic efficiency

We first demonstrated the functionality of our flow-through system by monitoring NP and Y(II) under SI and DI photoperiods (Figure 2). Results showed that while the irradiance regime did not significantly impact total daily O_2_ production (Student’s paired t-test; df = 3, t =-0.177, p = 0.871) (Figure 2B), dynamic changes in NP and Y(II) throughout the day could be detected, and were strongly shaped by the irradiance regime (Figure 2). Though NP was highest during midday hours in both treatments, the amplitude and rate of change in NP throughout the day was higher under DI, while remaining more stable, especially during the midday hours, under SI (Figure 2A).

PAM fluorometry showed that patterns of Y(II) also differed between irradiance regimes (Figure 2C, D). Under SI, daytime Y(II) was highest during the mid-to late morning, and there was a statistically significant negative linear relationship between Y(II) and time throughout the day (linear regression; p = 0.004, R^2^ = 0.217; Figure 2C). Under DI, Y(II) also changed throughout the day, but followed a different pattern inverse to irradiance (polynomial regression; p < 0.001, R^2^ = 0.433), as Y(II) was highest in the morning and evening, and lowest at midday (Figure 2D).

In contrast to the difference in patterns of Y(II) during the day (Figure 2C, D), dark-adapted Fv/Fm measured at night did not differ between treatments (LMM, F(1, 4) = 1.56, p = 0.279) and remained stable over time (linear regression; R^2^ = 0.042, p = 0.579 for SI; R^2^ = 0.049, p = 0.660 for DI).

### Experiment 2 - Daytime respiration estimates

Peak respiration rates detected during 6-minute periods of darkness introduced at midday showed that daytime R was 25% higher (mean ± SE of 0.109 ± 0.026 μmol O_2_ consumed s^-1^ g host protein^-1^) than nighttime R (mean ± SE of 0.087 ± 0.024 μmol O_2_ consumed s^-1^ g host protein^-1^) (Figure 3B; Student’s paired t-test; df = 4, t = 2.79, p = 0.049). Notably, light-enhanced dark R decreased towards the end of the 6 min measurement window, indicating a return towards nighttime rates (Figure 3D).

**Figure 3:**
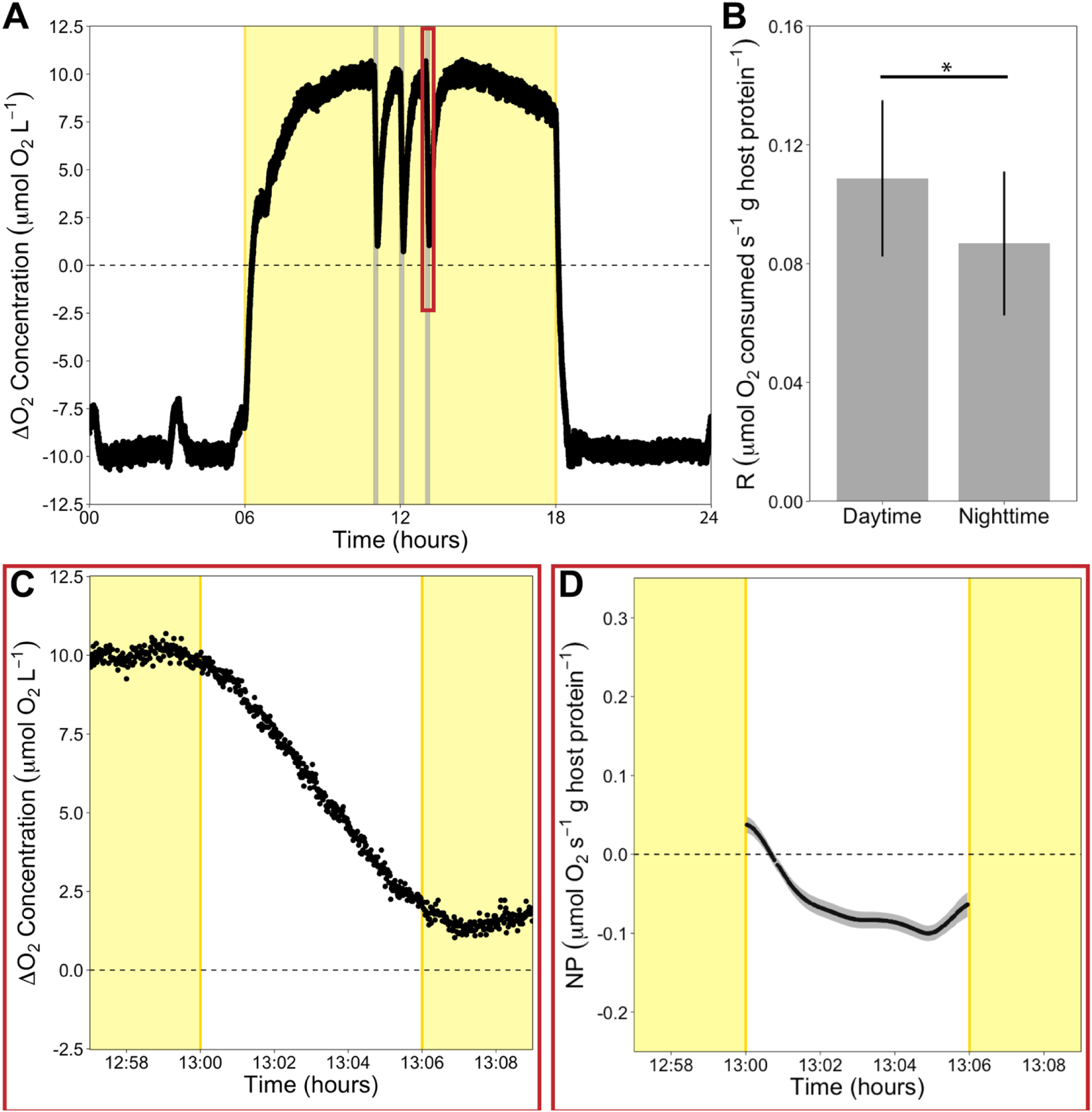
Quantification of daytime respiration. (A) Difference between outflow and inflow O2 concentration monitored continuously over 24h with SI photoperiod. Light-periods are highlighted in yellow, while grey lines mark 6-minute periods of darkness during the day, during which R was calculated (Equation 1). (B) Comparison of mean (± SE) daytime and nighttime respiration rates of photosymbiotic *Exaiptasia diaphana* CC7 holobionts (n = 5). (C) A closer view of the rapid change in outflow [O2] during a 6-minute period of darkness during the day (the red box indicated in A). (D) Daytime respiration was estimated using the minimum NP achieved during 6-minute periods of darkness during the day; mean (± SE) of all individuals (n = 5) across each period of darkness. SE = standard error; ΔO2 concentration = [O2]out-[O2]in. Asterisks above each bar indicate statistically significant differences.

### Experiment 3 - Comparison between closed and flow-through systems

Flow-through incubations reliably allowed quantification of NP and R in *E. diaphana* over full diurnal cycles at high temporal resolution. By comparing these integrated rates to extrapolated daily rates derived from closed-chamber incubations, we found that estimates derived from closed-chamber incubations significantly underestimate O_2_ production over a 12-hour photoperiod (Student’s two sample t-test, df = 7, t = 8.66, p < 0.001; Figure 4A), as well as O_2_ consumption over a 12-hour night (Student’s two sample t-test; df = 7, t = 3.07, p = 0.018; Figure 4B). Additionally, estimates of total O_2_ production over 24-hours in the flow-through system showed that *E. diaphana* holobionts were net productive (total daily O_2_ production of 1566 ± 2307 μmol O_2_ g host protein^-1^), while estimates from the closed system showed that they were net respiring (total daily O_2_ production of-1165 ± 340 μmol O_2_ g host protein^-1^) (Figure 4C). While the difference in total 24-hour O_2_ production was not statistically significant (Student’s two sample t-test; df = 7, t =-1.32, p = 0.227), daily P/R ratios showed P/R was 150% higher in the flow-through system (mean ± SE P/R of 1.38 ± 0.35) compared to the closed system (mean ± SE P/R of 0.55 ± 0.09) (Student’s two sample t-test; df = 7, t =-2.49, p = 0.042) (Figure 4D). This finding suggests significant underestimation of primary productivity when extrapolating from short-term, closed incubations, and may have implications for ecosystem-level contributions of benthic producers.

**Figure 4:**
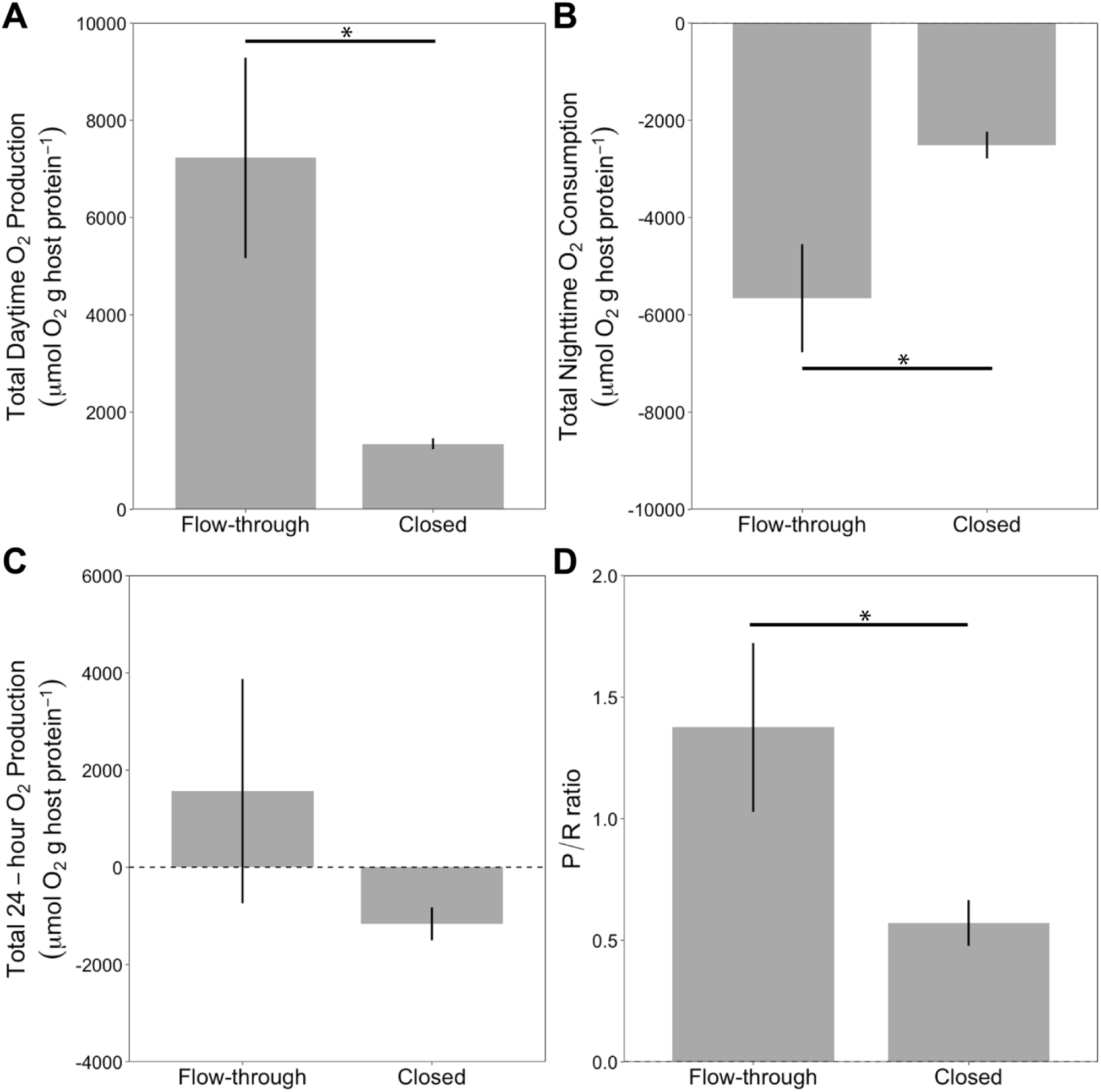
Comparisons of total O2 production and consumption in flow-through (n = 4) vs. closed systems (n = 5) with 12h:12h light/darkness cycle and SI photoperiod. Mean (± SE) (A) of net O2 production and (B) consumption over a 12-hour photoperiod and darkness, respectively. (C) Mean (± SE) total O2 production over a 24-hour period with 12h:12h light/dark cycle. (D) P/R ratios over the daily cycle; a value over 1 indicates net production of O2 over a daily cycle. Asterisks above bars indicate statistically significant differences.

### Experiment 4 - Long-term NP monitoring in flow-through

Long-term monitoring of NP showed that our photosymbiotic holobionts tolerated the conditions within the flow-through system for two weeks with no signs of stress of physiological impairment. Upon collection at the end of the experiment, the anemone appeared healthy, though there was mild algal growth visible; the chamber and water within were still transparent, but with a slight yellow tint from algal growth on the chamber walls that was not present in the negative control chamber.

Total nighttime O_2_ consumption changed little during the experiment, especially the first week, with night-to-night changes in consumption less than 8%, oscillating around a mean ± SE of 4786 ± 53 μmol O_2_ g host protein^-1^ consumed each night, and only a 4% difference between consumption on the first night and the seventh night (Figure 5; Table 1). The largest increase (12%) in nighttime O_2_ consumption occurred on night eleven, and was likely connected to the onset of mild algal growth within the chamber, given the trend in daytime O_2_ production on that same day (see below).

**Figure 5:**
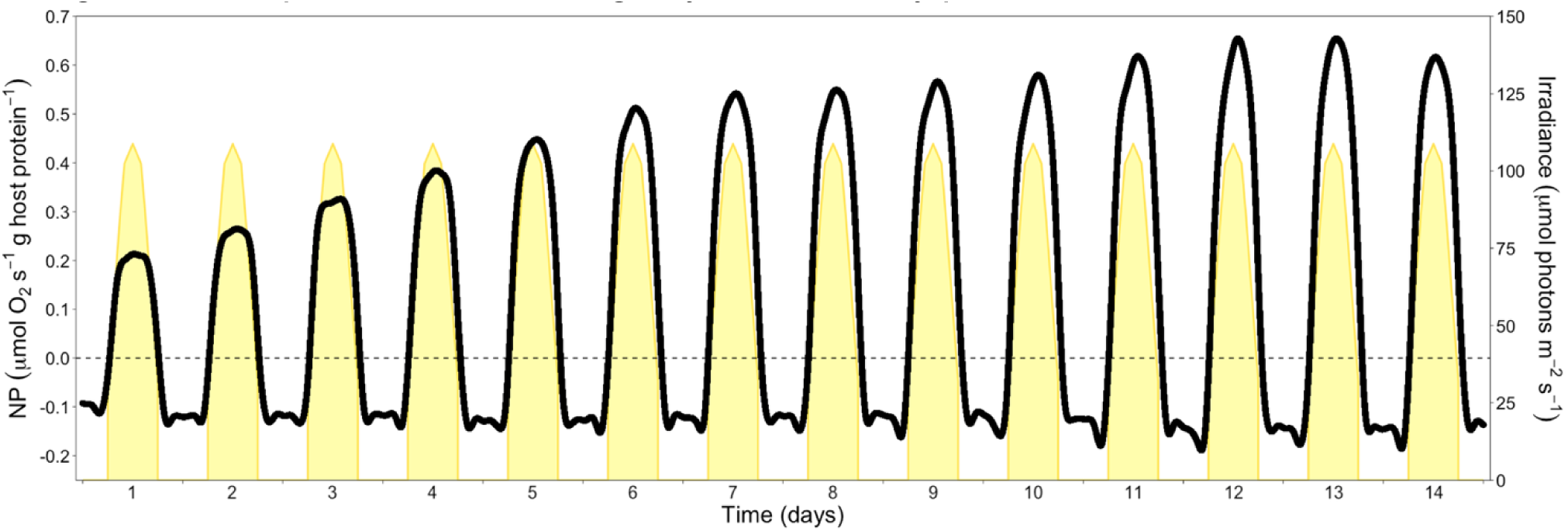
Continuous net productivity (NP) monitoring of an individual *Exaiptasia diaphana* holobiont within the flow-through system over two weeks. Light-periods are highlighted in yellow.

**Figure 6:**
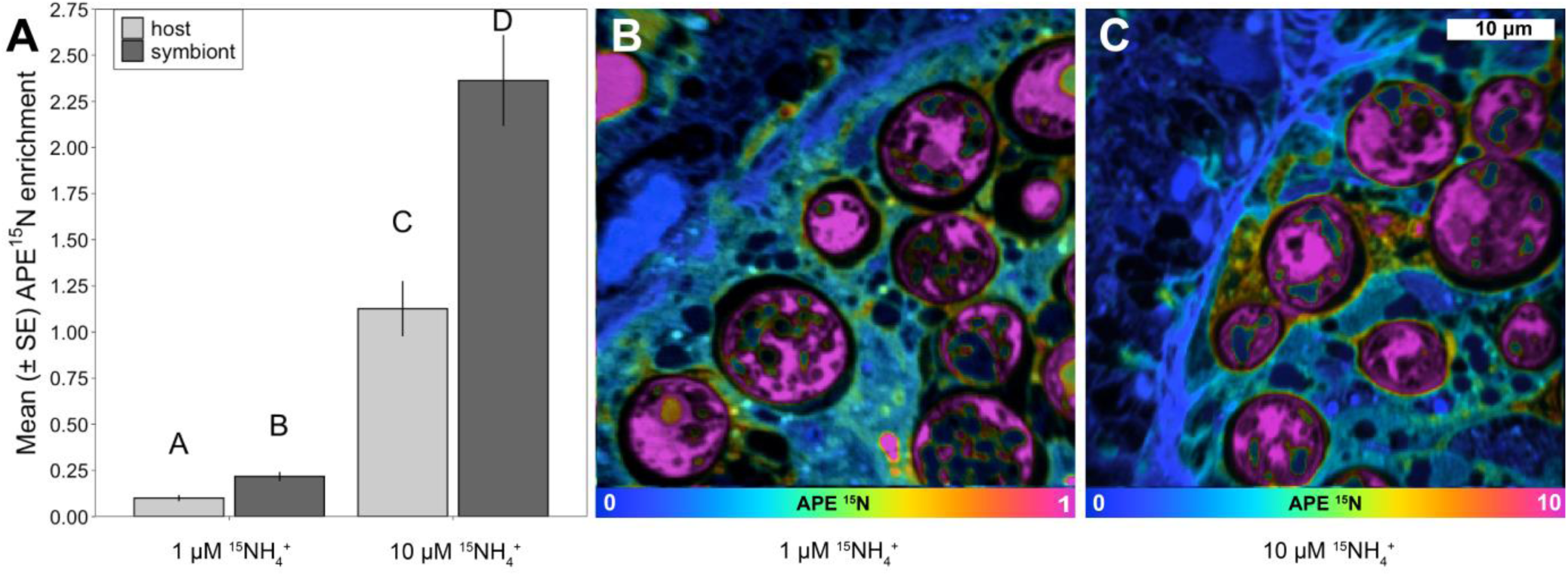
^15^N assimilation in the photosymbiotic *Exaiptasia diaphana* CC7 holobiont. (A) Comparison of mean (± SE) host and symbiont bulk tissue (n = 3 per host/symbiont in 10 μM treatment and n = 4 in 1 μM treatment) enrichment in ^15^N between treatments measured using IRMS. Different letters above each bar indicate statistically significant differences (p < 0.05). (B, C) NanoSIMS images showing ^15^N enrichment of host tissue and algal symbiont cells within 45 x 45 μm areas after a 24-hour incubation in the flow-through system with seawater containing either 1 μM ^15^NH ^+^ (B) or 10 μM ^15^NH ^+^ (C). Color scale bars represent enrichment in units of atom percent excess (APE), where pink is more highly enriched and blue is unenriched.

**Table 1:** Total net O2 production (during the 12-hour photoperiod) and total O2 consumption (during the night) are shown for each day of a 14-day long monitoring experiment, along with the % change from the previous day.

| **Day** | **Total daytime net O<sub>2</sub> production (µmol O<sub>2</sub> g host protein<sup>-1</sup>)** | **% Change from previous day** | **Total nighttime O<sub>2</sub> consumption (µmol O<sub>2</sub> g host protein<sup>-1</sup>)** | **% Change from previous day** |
|---|---|---|---|---|
| 1 | 6909 | N/A | 4790 | N/A |
| 2 | 8768 | + 27% | 4749 | - 1% |
| 3 | 11010 | + 26% | 4647 | - 2% |
| 4 | 12691 | + 15% | 5015 | + 8% |
| 5 | 14812 | + 17% | 4931 | - 2% |
| 6 | 16648 | + 12% | 4735 | - 4% |
| 7 | 17799 | + 7% | 4638 | - 2% |
| 8 | 17995 | + 1% | 4706 | + 1% |
| 9 | 18301 | + 2% | 4753 | + 1% |
| 10 | 18812 | + 3% | 4980 | + 5% |
| 11 | 19923 | + 6% | 5596 | + 12% |
| 12 | 20499 | + 3% | 5494 | - 2% |
| 13 | 20825 | + 2% | 5562 | + 1% |
| 14 | 19617 | - 6% | 5290 | - 5% |

In contrast to nighttime O_2_ consumption, total O_2_ production during the day increased greatly during the first 6 days, before plateauing after day 7 (Figure 5), likely due to photoacclimation within this period, as the individual used here had been maintained in SI but was subjected to DI during the experiment. Indeed, total O_2_ production increased by greater than 10% every day for the first 6 days, and an additional 7% on day 7, such that total O_2_ production on day 7 (17799 μmol O_2_ g host protein^-1^) was 157% higher than on day 1 (6909 μmol O_2_ g host protein^-1^) (Table 1). However, there was only a 6% change in O_2_ production between day 7 and day 10 (Table 1), and only 10% difference between O_2_ production on day 14 and day 7 (Figure 5; Table 1). A mild increase in daytime O_2_ production on day 11 is worth noting, as it coincided with the increase in O_2_ consumption at night, further suggesting onset of mild algal growth within the chamber at this point.

The stability of holobiont physiological performance shown here demonstrates that experiments lasting at least 10 days may be conducted using the flow-through system, though our ability to accurately estimate holobiont metabolism may be affected by algal overgrowth in experiments extending beyond a 10-day period.

### Experiment 5 - Flow through incubations permit ^15^NH_4_^+^ isotope labeling at low concentrations

IRMS results revealed significantly (11-fold) higher ^15^N enrichment in the high [^15^NH_4_^+^] treatment compared to the low [^15^NH_4_^+^] treatment (ANOVA; df = 1, F = 364.33, p < 0.001) (Table 2). Additionally, ^15^N enrichment of symbiont material was significantly higher than that of the host (ANOVA; df = 1, F = 37.59, p < 0.001) in both treatments (Table 2), though there was no interaction effect of nutrient concentration and partner identity (ANOVA; df = 1, F = 0.03, p = 0.869).

**Table 2:** Mean ±SE ^15^N enrichment (in APE) of host and symbiont fractions of *Exaiptasia diaphana* holobionts. N = 4 for each treatment of each method.

| **Method** | **[<sup>15</sup>NH<sub>4</sub><sup>+</sup>]** | **Mean Host** | **SE Host** | **Mean Symbiont** | **SE Symbiont** |
|---|---|---|---|---|---|
| IRMS (bulk tissue) | 10 | 1.126 | 0.149 | 2.363 | 0.247 |
| IRMS (bulk tissue) | 1 | 0.101 | 0.017 | 0.217 | 0.025 |
| NanoSIMS (cellular scale) | 10 | 3.248 | 0.713 | 13.358 | 1.921 |
| NanoSIMS (cellular scale) | 1 | 0.208 | 0.058 | 0.914 | 0.294 |

In addition to IRMS findings, data from host and symbiont ROIs in NanoSIMS images showed that our flow-through system avoided nutrient depletion in the low [^15^NH ^+^] experiment, with sufficient labelling to measure enrichment at the subcellular level. Statistical analysis of ^15^N enrichment measured within individual ROI showed similar results to IRMS analysis, with a significant effect of treatment (LMM; F(1, 5) = 37.51, p = 0.002) and host/symbiont identity (LMM; F(1, 71) = 124.37, p < 0.001), but not the interaction of treatment and host/symbiont identity (LMM; F(1, 71) = 0.56, p = 0.455) on ^15^N enrichment (Table 2).

## Discussion

Coastal and estuarine ecosystems are characterized by dynamic environmental conditions, yet our ability to quantify physiological performance of benthic organisms in these environments has long been constrained by well-known limitations of conventional laboratory assays. To address this, we developed a novel experimental flow-through system that builds upon established laboratory techniques to allow the introduction of environmental variability in a controlled setting. By enabling continuous monitoring of physiological metrics under controlled yet dynamic laboratory conditions over days to weeks, the modular design of this platform provides a robust foundation for addressing complex biological responses ranging from symbiotic interactions in aquatic holobionts to global change impacts on key foundational species.

We demonstrated that our flow-through system provides advantages over traditional closed-chamber incubations, which permit efficient short-term measurements with high replication (Liu et al., 2020; Lyndby et al., 2020; Sawall et al., 2022). Despite their efficiency, conventional experiments must balance chamber volume against metabolic activity to prevent accumulation of metabolic waste products or depletion of O_2_ and nutrients (Camp et al., 2015), which may have feedback effects on physiological performance and complicate interpretation (Camp et al., 2015; Liu et al., 2020; Sawall et al., 2022). Additionally, reliance on short incubation periods necessitates extrapolation to longer timescales. As shown here, this approach might significantly underestimate productivity and respiration and result in biased P/R ratios. In contrast, our flow-through system mitigates these issues by maintaining a constant supply of new seawater, stabilizing O_2_ and nutrient availability throughout incubations that can consequently last longer than one week.

By allowing for continuous observations over days to weeks, the flow-through system captures organismal responses to environmental fluctuations as they occur. This contrasts to conventional methods for understanding of organismal responses to environmental change, utilizing snapshots of physiological performance before and after exposure to a change in environmental conditions (Godinot et al., 2011; Rädecker et al., 2017; Röser et al., 2024). Thus, continuous monitoring is vital for improving our understanding of organismal responses to environmental change and how these responses may affect species’ ecosystem-level contributions (Sawall et al., 2025). We demonstrated that methodology has a strong effect on estimation of O_2_ flux on a daily scale, with important consequences for the interpretation of holobiont physiology. P:R ratios – commonly used as an indicator of the metabolic balance of the holobiont – estimated in the flow-through system indicated net O_2_ production on a daily scale, while the opposite was indicated according to closed incubations. Therefore, the two experimental approaches could lead to differing conclusions about the trophic status and energetic balance of the organism. The higher rates measured under continuous flow may result from a longer recovery period for individuals after being manipulated prior to experimentation, the maintenance of more stable environmental conditions around the anemones (avoiding large changes in O_2_ and CO_2_ conditions following photosynthesis and respiration) during the experiment, or a combination of these two factors. Studies on cnidarians have shown that increased flow can enhance both respiration and photosynthesis, partly by improving gas exchange and CO₂ availability (Osinga et al., 2017). Thus, in addition to more stable conditions within the chamber during experimentation, an improved gas exchange may contribute to the higher rates of O_2_ production and consumption measured during the day and night, respectively, in the flow-through system compared to closed incubations.

Beyond improving the precision of metabolic estimates, the responsiveness of the flow-through system permits quantification of daytime R. Conventional respirometry methods typically rely on the measurement of nighttime R and the assumption that R remains constant over the full 24-hour cycle, though it is widely accepted that this likely underestimates respiratory demand during the daytime and, consequently, leads to an erroneous calculation of GP (Hudspith et al., 2022; Kühl et al., 1996; Owen et al., 2021; Sawall et al., 2025). Our results confirm this postulate. By incorporating brief periods (here 6 minutes) of darkness during the photoperiod, our flow-through system enables repeated quantification of daytime R. While these estimates likely still underestimate rates compared to microsensing techniques (Kühl et al., 1995, 1996), our approach balances the need for high temporal resolution with the need for holobiont-scale characterization.

The ability to integrate O_2_ flux data with real-time PAM fluorometry offers a mechanistic lens into the interplay between productivity and photophysiology. By linking changes in primary productivity to specific shifts in effective quantum yield (Y(II)), the system permits differentiating between the metabolic consequences of environmental fluctuations and underlying photo-acclimation processes. While photoinhibition is commonly associated with severe light stress, our results highlight the system’s capacity to detect even a mild dynamic down-regulation of PSII photochemistry at low light intensities (Schreiber et al., 1995), and this in the context of measured NP. Additionally, though metrics of Y(II) were not collected during the long-term monitoring experiment, trends in daytime O_2_ production over two weeks suggest that within approximately one week photosymbionts of *E. diaphana* become photoacclimated to a new irradiance regime, which can serve as a guide when exposing organisms to new environmental conditions.

The system’s constant flow design also enhances our ability to conduct isotopic labeling studies by allowing use of environmentally realistic nutrient concentrations, while still reaching enrichment levels that are easily measured via both IRMS and NanoSIMS. This improves upon past methods by avoiding excessive nutrient loads that may exacerbate adverse effects brought on by other environmental stressors (Rädecker et al., 2023; Zhao et al., 2021).

## Future directions

Physiological processes within marine holobionts directly affect the metabolites they exchange with the surrounding environment. For example, heat stress may enhance the release of metabolic waste products such as CO_2_ or NH ^+^ from the holobiont or trigger oxidative stress responses resulting in the release of reactive oxygen species (Dong & Zhang, 2016; Madeira et al., 2018). We are currently working on developing a 3D-printable autosampler for the flow-through system to collect outflow seawater, allowing for simultaneous measurements of holobiont nutrient flux and water chemistry at high temporal resolution. Such monitoring of effluent water chemistry will permit detection of nutrient flux within the system, allowing evaluation of whether a study organism serves as a source or sink of key nutrients, such as HCO ^-^ or NH ^+^. Detection of changes in nutrient release by the holobiont can be evaluated in the context of other data collected, including isotopic labelling or O_2_ monitoring, thus providing a thorough assessment of holobiont physiological performance.

Additionally, a reactive oxygen species (ROS) sensor is currently being developed based on a microsensor already tested in other workflows (Ousley et al., 2022) and may allow for estimation of holobiont H_2_O_2_ release. The incorporation of this sensor has been principally driven by interest in ROS production within cnidarian holobionts, as production of ROS has long been viewed as a hallmark signal in the breakdown of the cnidarian-algal symbiosis during heat stress (Dunn et al., 2012; Gardner et al., 2017; Lesser, 1996; Roberty et al., 2015). However, its application may also be used to resolve stress responses and signaling cascades in other biological systems.

## Conclusion

Our novel flow-through system builds upon existing methods to provide a robust approach for simultaneous monitoring of a variety of biological activities. A constant supply of well-oxygenated seawater with standardized water chemistry allows incubations that last days (or potentially weeks to months) at a time. Thus, its implementation within incubators that give control over key environmental factors provides a means for the recreation of marine heatwaves and other stressful events in a highly controlled setting. Use of this novel technique and its adaptation for other species and methodologies, such as ecotoxicology, will improve our mechanistic understanding of the effects of a variety of environmental stressors that are commonly experienced in the Anthropocene. By doing so, researchers and stakeholders may be able to better inform key decisions regarding conservation and management of vulnerable coastal ecosystems.

## Acknowledgements

We thank Jules Tresca for his diligent caretaking of cultured anemones, as well as with sample collection and processing. Additionally, we thank Dr. Laszlo Kocsis and Dr. Thomas Kuhn for providing preliminary IRMS analyses, as well as Cristina Martin Olmos and Stéphane Escrig for their assistance in obtaining SEM and NanoSIMS images. We also thank Nele Eichhorn for creating the illustration in Figure 1.

## Data Availability

All data and R scripts associated with this publication are freely available at 10.5281/zenodo.24377869.

## Author contributions

HJ, CP, AM and NR conceived and designed the study. HJ performed the experiments and analysed the data. CF-P and RG contributed to sample analysis. HJ wrote the first draft of the manuscript, and all authors contributed critically to its revision and gave final approval for publication.

